# Repeated self-administered EEG recordings enable longitudinal measurement of brain activity at home

**DOI:** 10.64898/2026.11.12.766143

**Authors:** Dirk van Moorselaar, Ruchella Kock, Jan Theeuwes

**Affiliations:** Axite, Leiden, the Netherlands; Department of Experimental and Applied Psychology, Vrije Universiteit Amsterdam, the Netherlands; Institute of Brain and Behaviour Amsterdam (iBBA), the Netherlands; William James Center for Research, ISPA-Instituto Universitario, Lisbon, Portugal

## Abstract

Mobile electroencephalography (EEG) offers new opportunities to investigate brain activity outside the laboratory, but the quality and reliability of self-administered recordings remain important concerns. Here, we evaluated whether repeated at-home recordings using a four-channel consumer EEG device could provide meaningful electrophysiological measurements and characterized the stability of these measures across sessions. Five participants completed repeated recordings across multiple days, including resting-state measurements and an auditory oddball task performed with and without concurrent cognitive load. The recordings captured established spectral signatures of physiological state and task demands, including increased alpha activity during eye closure and changes in theta activity associated with cognitive load. Auditory stimulation elicited clear event-related responses, while exploratory multivariate decoding distinguished standard from target tones across recording sessions. Measurement stability varied across electrophysiological features, and aggregating repeated sessions improved the consistency of individual-level estimates. For example, split-half reliability of the auditory N1-like response increased from r = .22 with one session to r = .76 with seven sessions per half. By demonstrating the feasibility of repeated self-administered EEG measurements, this study provides a step toward investigating how brain activity varies both between individuals and within individuals over time in everyday environments.

## Introduction

Electroencephalography (EEG) offers millisecond-level temporal resolution of neural activity, making it a central tool for studying cognitive dynamics and for diagnosing and monitoring neurological conditions (Luck, 2014; Michel & Murray, 2012). Conventional EEG systems, however, typically require gel-based electrode preparation, dedicated recording equipment, and trained personnel, limiting their use outside supervised laboratory or clinical settings. Mobile EEG systems, encompassing wearable devices with varying electrode configurations and channel densities, offer a more accessible alternative that can be deployed outside the laboratory with relatively little training. Although low density consumer-grade systems achieve this accessibility at the cost of reduced spatial coverage and potentially greater susceptibility to recording noise and artefacts, previous studies have demonstrated that established EEG phenomena, including spectral dynamics and event-related potentials, can be measured using these devices (Cannard et al., 2021; Krigolson et al., 2017; Sabio et al., 2024).

Despite this promise, validation research using mobile and consumer-grade EEG has primarily focused on establishing signal quality and demonstrating that these devices can capture established neural phenomena (Sabio et al., 2024). A complementary question is whether the resulting measures are sufficiently reliable at an individual level (Lau-Zhu et al., 2019). This question is particularly important because some of the most compelling opportunities afforded by mobile EEG arise from its accessibility: repeatedly tracking neural measures within the same individual over days or weeks, and examining stable differences in these measures between individuals. Realizing these opportunities requires moving beyond demonstrating that an EEG effect can be detected at the group level to establishing whether it provides a consistent and reliable measurement of an individual.

What constitutes a reliable measurement, however, depends on how that measurement is intended to be used (Lopez et al., 2023). For individual-differences research, repeated measurements should preserve stable differences between individuals, such that individuals who consistently show higher values on a given EEG marker can be distinguished from those who show lower values. For longitudinal applications, a complementary concern is whether repeated measurements within an individual are sufficiently stable to support the interpretation of changes over time. Although both applications require that the signal of interest can first be detected, they place different demands on the consistency of the resulting measurements. A measure that reliably distinguishes between individuals is therefore not necessarily sufficiently stable to track changes within an individual over time. Moreover, demonstrating an expected EEG effect at the group level does not establish either form of reliability. Indeed, experimental effects can be highly robust at the group level while showing limited reliability for distinguishing between individuals, particularly when experimental paradigms are designed to minimize between-participant variability (Hedge et al., 2018).

Research using conventional laboratory EEG has demonstrated that spectral measures can provide reliable individual-level estimates across repeated recordings. Resting-state spectral power generally shows good-to-excellent test–retest reliability, particularly for alpha-band activity (Lopez et al., 2023). For example, Näpflin et al. (2007) demonstrated that individual characteristics of resting-state EEG spectra remained sufficiently stable to distinguish participants across recording sessions separated by more than one year, while other studies have reported high test–retest reliability for theta, alpha, and beta power during both resting-state and cognitive-task recordings (Ip et al., 2018; McEvoy et al., 2000). Importantly, reliability is not restricted to resting-state activity. Ding et al. (2022) found that the reproducibility of spectral measures differed across recording conditions, with eyes-closed rest showing higher reproducibility than eyes-open rest at the sensor level, while mental arithmetic produced comparatively high reproducibility for several frequency bands. Furthermore, recent work indicates that derived spectral indices, including ratios between frequency bands, can also exhibit high test–retest reliability (Chang, 2025). Together, these studies demonstrate that reliable spectral measurements are not limited to resting-state power, but can also extend to task-related activity and derived spectral indices.

Event-related potentials (ERPs) can also provide reliable individual-level measurements, although reported test–retest reliability varies considerably across components and quantification approaches (Lopez et al., 2023). For example, Cassidy et al. (2012) reported moderate-to-strong reliability across several ERP components measured one month apart, while studies using auditory oddball paradigms have demonstrated reliable individual differences in P300 amplitude across repeated recordings (Debener et al., 2002; Segalowitz & Barnes, 1993). Importantly, reliability also depends on methodological factors, including the number of trials contributing to an ERP average and the selection of recording electrodes (Huffmeijer et al., 2014). At the same time, repeated exposure to auditory oddball tasks can produce systematic changes in ERP amplitudes, including reductions in P300 amplitude across sessions (Debener et al., 2002; Kinoshita et al., 1996). Together, these findings indicate that ERPs can provide reliable individual-level measurements under conventional laboratory conditions, although their reliability depends on the component, measurement approach, and potential systematic changes associated with repeated testing.

To the best of our knowledge, the reliability of both spectral and event-related measures obtained using low-density consumer EEG has not been systematically characterized across densely repeated recordings within multiple individuals. Nevertheless, preliminary evidence suggests that reliable measurements are possible with consumer and wearable EEG systems, although findings vary across devices, neural measures, and recording conditions (Sabio et al., 2024). For example, Rogers et al. (2016) reported good test–retest reliability for resting-state spectral measures obtained using a single channel consumer EEG device. Using a combination of scalp and ear-EEG, Mikkelsen et al. (2021) compared repeated recordings collected in the laboratory and at home, finding that between-individual differences generally exceeded variability attributable to recording location, although some measures showed greater unexplained variability at home. More recently, Barbey et al. (2022) demonstrated that repeated, self-administered EEG recordings could be collected successfully at home using a higher density wearable system. By quantifying the precision of ERP amplitude estimates, they further showed that aggregating trials across multiple home recordings could yield measurement precision comparable to conventional laboratory EEG. Together, these studies provide encouraging evidence for the feasibility and measurement quality of EEG recordings beyond conventional laboratory settings. However, the consistency of both spectral and event-related measures obtained using low-density consumer EEG, and the extent to which this consistency improves with session aggregation, remain less well established.

To investigate whether low-density consumer EEG can provide reliable individual-level measurements across repeated recordings, we collected densely sampled EEG data from the same individuals using the four-channel Muse S system. Muse provides an informative test case because it is a widely used and accessible consumer EEG system for which established spectral and event-related effects have previously been demonstrated (Kleeva et al., 2024; Krigolson et al., 2017). In each session, participants completed resting-state recordings with eyes open and closed, providing a well-established manipulation of alpha activity (Barry et al., 2007), followed by an auditory oddball task performed either alone or concurrently with mental arithmetic to increase cognitive load. This design allowed us to investigate several classes of EEG measures: resting-state spectral activity, task-related spectral dynamics associated with cognitive load, including theta- and alpha-band activity and their ratio (Chikhi et al., 2022), and event-related responses to auditory targets (Polich, 2007). We first examined whether the expected EEG effects could be detected at the individual level, before assessing the extent to which the resulting measures preserved stable differences between individuals and remained consistent within individuals across repeated recordings. Finally, the dense sampling allowed us to investigate how measurement reliability changes when information is aggregated across sessions, providing an estimate of how much data may be required to obtain reliable individual-level measurements using low-density consumer EEG.

## Methods

### Participants

Participants were recruited from individuals who had previously taken part in research conducted by Axite and had indicated that they were willing to be contacted about follow-up studies. Eligible participants were healthy adults aged 18 years or older, proficient in English, with no history of neurological or psychiatric disorders. Seven participants entered the study, of whom five contributed sufficient usable EEG data for at least one experimental contrast. Participant inclusion required at least 17 paired recording sessions meeting the initial data-quality criteria for the relevant contrast. This threshold was selected to retain a participant who provided good-quality recordings but did not complete the full recording protocol. Five participants were included in the single- versus dual-task analyses, and four in the eyes-open versus eyes-closed analyses. Subsequent channel- and region-specific quality-control exclusions reduced the number of usable sessions for some participants but did not change the participant sample. The final sample comprised five participants (mean age = approaximately 29 years, range = 23–43; 3 female). All participants provided written informed consent before participation. The study was approved by the Ethical Review Committee of the Faculty of Behavioural and Movement Sciences of the Vrije Universiteit Amsterdam.

### Apparatus and software

EEG was recorded continuously at a sampling rate of 256 Hz using a four-channel Muse S Athena headband (InteraXon, Toronto, Canada), with electrodes positioned at AF7, AF8, TP9, and TP10 and referenced to Fpz. The headband additionally provided onboard accelerometer and photoplethysmography (PPG) data. EEG acquisition and the behavioural task were run on an Android smartphone, using either the participant’s own device or a study-provided device when necessary. The behavioural task was implemented in OpenSesame 4 (Mathôt et al., 2012) using the OSWeb runtime and delivered through JATOS (Lange et al., 2015). EEG acquisition was handled by NeuroJournal (Axite B.V.), a purpose-built Android application that communicates with the Muse headband via Bluetooth and provides in-app onboarding and real-time signal-quality feedback to support self-administered recordings. EEG preprocessing was performed using Axite’s custom Python based processing software, which incorporates MNE-Python (Gramfort et al., 2014). Subsequent statistical analyses were performed using custom Python scripts.

### Design and stimuli

Each session included two experimental manipulations designed to elicit established EEG responses. First, resting-state EEG was recorded during alternating eyes-open and eyes-closed periods to examine changes in alpha-band activity associated with eye closure (Barry et al., 2007). Second, participants completed an auditory oddball task under single- and dual-task conditions to investigate event-related responses to auditory targets and spectral changes associated with increased cognitive load. Each oddball block consisted of 300 tones, comprising 500-Hz standards and 1000-Hz targets, with an intended target probability of approximately 20%. The block length was chosen to provide approximately 60 target trials per condition while keeping session duration suitable for repeated self-administered recordings. Each block began with 5–8 standard tones, and successive targets were separated by at least two standards. Beyond these constraints, target occurrence was determined probabilistically, with the assignment probability adjusted to maintain an expected overall target proportion of approximately 20%. Each tone lasted 60 ms with cosine-ramped onset and offset, and stimulus-onset asynchronies were sampled uniformly between 800 and 1200 ms. In the single-task condition, participants attended to the tone stream and had to count the number of target tones. They provided an approximate estimate of the number of targets at the end of each block. In the dual-task condition, participants additionally performed silent serial subtraction, a cognitively demanding task previously associated with changes in EEG theta activity (Chikhi et al., 2022; Sammer et al., 2007). Participants repeatedly subtracted a fixed number (17 by default, adjusted across sessions when necessary to maintain task difficulty) from a starting value randomly selected between 1001 and 1029 throughout the block, and reported both their target estimate and final subtraction total at the end of the block. The order of the single- and dual-task conditions, as well as the initial eyes-open/eyes-closed state, was randomized across sessions.

### Protocol

Participants were asked to complete 28 recording sessions of approximately 15 minutes each, with a target frequency of two sessions per day. The twice-daily schedule was intended to enable exploratory comparisons between morning and evening recordings, motivated by the possibility that EEG measures might vary with time of day and associated differences in alertness or fatigue. Following an initial supervised training session, recordings were self-administered at home, with additional supervised sessions provided when needed. During training, participants received guidance on fitting the headband and optimizing signal quality. They were also instructed to adjust the difficulty of the serial-subtraction task across sessions when necessary to maintain an appropriate level of challenge, using 17 as the default subtrahend. Although participants were encouraged to complete two sessions per day, the self-administered nature of the protocol allowed flexibility in scheduling, resulting in variation in recording frequency and overall study duration between participants. Among the five participants included in the present analyses, recordings spanned a median of 36.7 days (range: 22.6–50.4 days). The median interval between successive sessions was 0.55 days across participants (range of participant-specific medians: 0.50–1.23 days), although individual intervals were occasionally substantially longer, reaching up to 20.2 days.

At the start of each session, participants connected the headband to the NeuroJournal app and confirmed adequate signal quality before beginning the recording. Each session began with five visually cued blinks followed by five taps on the headband, which provided identifiable EEG and accelerometer responses for offline verification of temporal correspondence between the behavioural and physiological recordings. Participants then completed four alternating 60-s resting-state periods (two eyes-open and two eyes-closed), followed by the single- and dual-task auditory oddball blocks. Each oddball block was preceded by three cued screen-touch responses to support synchronization of behavioural event timestamps. The session concluded with a second calibration sequence of five cued blinks and five headband taps. Participants could subsequently add free-text notes describing events that might have affected the recording, such as signal-quality warnings or periods of distraction.

### Preprocessing

Raw EEG was linearly interpolated onto a uniform 256-Hz sampling grid and subsequently band-pass filtered between 1 and 40 Hz using a zero-phase, Hamming-windowed finite impulse response (FIR) filter. No additional re-referencing was performed, preserving the headband’s native Fpz reference. For ERP analyses, the filtered signals underwent an additional 30-Hz low-pass filtering step before epoch extraction. All preprocessing was performed using Axite’s proprietary Python-based processing pipeline, incorporating MNE-Python (Gramfort et al., 2014).

Signal contamination was identified using two complementary procedures. First, channel-specific artifacts were detected in non-overlapping 250-ms windows using features characterizing signal amplitude, flatness, temporal variability, and high-frequency activity. Detection thresholds were determined relative to locally adaptive reference distributions, calculated using rolling 60-s medians and median absolute deviations (MADs). The same detection procedure and parameter settings were applied to all participants and sessions. Second, blinks were identified through template matching using participant-specific waveforms derived from calibration recordings. Detected blinks were masked across all four EEG channels. Both procedures generated sample-level artifact masks without modifying or reconstructing the underlying EEG signals.

Artifact masks were applied separately for spectral and event-related analyses. For spectral analyses, contaminated windows were excluded on a channel-specific basis, followed by additional rejection of windows with extreme spectral power, as detailed below. For ERP analyses, individual channel-specific epochs were excluded when more than 15% of their samples were masked, while valid channels from the same trial remained available for analysis. No interpolation or reconstruction was performed to replace excluded data.

To identify sustained channel abnormalities that could remain undetected by these procedures, we applied an additional quality-control step to the session-level spectral measurements. For each participant, channel, and recording segment, spectral power was compared across sessions using a median- and MAD-based outlier criterion. A channel was considered persistently abnormal within a session when its spectral measurements exceeded a modified z-score threshold of 5 in at least 50% of the session’s segments, with a minimum of four segments required for evaluation. Sessions containing such abnormalities were excluded from analyses of the affected recording region (frontal: AF7/AF8; temporal: TP9/TP10), while measurements from the unaffected region were retained. This region specific exclusion was applied consistently to spectral and ERP analyses. For multivariate decoding, which combined features from both regions, sessions were excluded entirely when either region failed quality control.

### Temporal alignment

EEG samples and behavioural events were aligned using a common Unix-time reference, with OpenSesame event timestamps converted to Unix time using recorded browser timing information. Temporal correspondence between these streams was evaluated using several complementary procedures. First, calibration sequences of visually cued blinks and headband taps, performed at the beginning and end of each session, provided identifiable EEG and accelerometer responses for assessing the consistency of event timing across recordings. Second, an independent bench test using phone-generated vibrations detected in the Muse accelerometer recordings yielded timing differences centred near zero (median = −2.1 ms, SD = 40.9 ms), supporting approximate cross-device synchronization in the tested configuration. Third, a touchscreen calibration procedure comparing OpenSesame timestamps with independently recorded phone-side events revealed a mean residual difference of 17.6 ms (SD = 5.0 ms) across 26 sessions, supporting the accuracy of the browser to-OpenSesame clock conversion. Because ERP analyses additionally depend on consistent stimulus presentation timing, auditory onset variability was assessed using external acoustic recordings of a shortened version of the oddball task. Across two recordings of 50 tones each, differences between acoustically measured and logged inter-onset intervals had standard deviations of 1.73 and 2.00 ms, respectively, indicating low variability in relative auditory presentation timing. Together, these controls support approximate temporal correspondence between the recorded streams and consistent relative stimulus timing in the tested configurations. However, because Muse timestamps reflect sample receipt rather than independently verified acquisition times, and the absolute delay between logged auditory events and physical sound onset was not established, absolute ERP component latencies should be interpreted with caution. No additional fixed temporal correction was applied to the auditory event timestamps.

### Spectral power

Spectral power was estimated from 2-s analysis windows with 50% overlap. Artifact rejection was performed independently for each channel, excluding windows in which more than 25% of samples were flagged as contaminated. Retained windows were analyzed using their original signal values. Power spectral density (PSD) was calculated for each retained window using Welch’s method, with 1-s Hann-windowed segments, 50% overlap, and a zero-padded Fourier transform yielding 0.25-Hz frequency-bin spacing. Windows with extreme broadband (4– 30 Hz) power were additionally excluded using a modified z-score threshold of 5, calculated relative to other windows from the same recording segment and channel. PSD estimates were then averaged across surviving windows, and absolute band power was calculated by trapezoidal integration within the theta (4–8 Hz), alpha (8–12 Hz), and beta (12–30 Hz) frequency bands. For resting-state conditions, band-power estimates were first calculated separately for each recording period and subsequently averaged across the two periods. Regional measures were obtained by averaging channel-level band power across AF7 and AF8 (frontal) or TP9 and TP10 (temporal), with available channels weighted equally. Averaging was intended to provide more robust regional estimates by reducing channel-specific variability, as the analyses focused on broad spectral changes rather than lateralized effects.

The theta-to-alpha ratio was calculated separately for individual analysis windows and channels before averaging across windows and electrodes. Because changes in conventional band-power measures may reflect both frequency-specific activity and shifts in the broadband aperiodic background, we additionally parameterized the power spectra using FOOOF (Donoghue, Haller, et al., 2020). This allowed us to assess whether experimentally induced changes in spectral power could be accounted for by changes in the aperiodic component rather than necessarily reflecting changes in narrowband activity. FOOOF was fitted separately to each session-, condition-, and region-level average spectrum over the 1–40 Hz range, using a fixed aperiodic model. The resulting fits provided estimates of the aperiodic offset and exponent, as well as the frequency, amplitude, and bandwidth of detected spectral peaks. To assess the contribution of aperiodic activity to experimental effects, we reconstructed the aperiodic component of each condition’s spectrum and calculated the corresponding band-power measures and theta-to-alpha ratio. The condition difference predicted by these aperiodic components was then subtracted from the observed condition difference, yielding a residual effect not accounted for by the fitted aperiodic background. These residual differences were averaged within participants before statistical testing across participants.

### Spectral statistical analysis and decoding

Effects of the experimental manipulations on spectral measures were assessed by comparing eyes-open with eyes-closed resting-state recordings and single with dual-task oddball recordings. For each measure, values were first averaged across sessions within each participant and condition, and condition differences were subsequently evaluated using two-sided paired-samples t-tests across participants. Analyses focused on alpha power for the eyes-open/eyes closed comparison and on theta power, alpha power, and the theta-to-alpha ratio for the single-/dual task comparison. To examine whether observed condition differences could be accounted for by changes in the aperiodic spectral background, additional one-sample t-tests were performed on the participant-level residual differences obtained after subtracting the corresponding aperiodic-predicted changes.

Sessions were eligible for a given experimental contrast when both conditions contained at least 20 artifact-free analysis windows and retained at least one valid measurement after spectral outlier rejection. Participants were included when at least 17 sessions met these criteria for the relevant contrast. Subsequent channel-level quality-control exclusions were applied separately to frontal and temporal measurements, without removing participants who had already met the initial inclusion criterion.

To assess whether spectral features could distinguish experimental conditions across recording sessions, we additionally performed within-participant multivariate decoding. Separate classifiers discriminated eyes-open from eyes-closed resting-state recordings and single-from dual-task oddball recordings using 18 spectral features encompassing band power, the theta-to-alpha ratio, spectral entropy, aperiodic parameters, oscillatory peak characteristics, and hemispheric asymmetry. Because the classifiers combined features from both recording regions, sessions failing the additional quality-control criterion in either region were excluded from both conditions of the corresponding contrast. Classification was performed using regularized logistic regression with leave-one-session-out cross-validation, such that models were trained on all but one session and evaluated on the held-out session. Features were standardized using parameters estimated exclusively from the training data within each fold. Observations with missing feature values were excluded rather than imputed, and all 18 features were retained without feature selection. Regularization strength was optimized within the training data using nested cross-validation. Statistical significance was assessed using 200 permutations in which condition labels were independently exchanged within sessions, preserving the paired structure of the data. For each permutation, leave-one-session-out classification was repeated using a fixed regularization strength determined from the original analysis, and classification accuracy was averaged across participants to construct a group-level null distribution.

### ERP analysis

For ERP analyses, the preprocessed continuous EEG was additionally low-pass filtered at 30 Hz using a zero-phase FIR filter and segmented into epochs from −200 to 800 ms relative to the logged tone onset. Baseline correction was applied separately to each trial and channel using the −200 to 0 ms prestimulus interval. Artifact masks from the continuous preprocessing pipeline were retained, and individual channel-by-trial epochs were excluded when more than 15% of their samples were marked as contaminated. Regional ERPs were obtained by averaging the available channels within the frontal (AF7, AF8) and temporal (TP9, TP10) regions. This averaging was intended to improve signal-to-noise ratio, as the experimental manipulations did not involve lateralized stimulation and visual inspection indicated comparable responses across hemispheres. Remaining trials were averaged separately for standard and target tones within each session and task condition. Participant-level waveforms were calculated by averaging across sessions, and group-level waveforms by averaging across participants. ERP amplitudes were expressed in the device’s native, uncalibrated units.

ERP quantification focused on three temporal measurement windows identified from participant-equal weighted grand-average waveforms. Each window was 100 ms wide and centered on a peak identified within a visually selected search interval. An N1-like negativity was identified at 375 ms in the stimulus collapsed waveform (300–450 ms search interval), yielding a measurement window of 325–425 ms. Two positive deflections were identified in the target-minus-standard difference waveform, collapsed equally across single- and dual-task conditions: an early positivity peaking at approximately 273 ms (150–350 ms search interval; 223–323 ms measurement window) and a late positivity peaking at approximately 641 ms (500–750 ms search interval; 591–691 ms measurement window). The late positivity served as the primary measure of target-related activity, with the early positivity treated as secondary. The measurement windows were selected based on visual inspection of the grand-average ERP waveforms to accommodate the observed component latencies, rather than being defined a priori from previous literature. Windows were fixed across participants, sessions, and conditions and were not subsequently optimized for reliability.

ERP amplitudes were quantified to address two questions. First, we examined whether ERP amplitudes differed between standard and target tones within the N1-like and late positivity windows. For these comparisons, responses were averaged across the single- and dual-task conditions, with both conditions weighted equally. Second, we examined whether cognitive load modulated target-related activity by comparing target-minus-standard amplitudes between the single- and dual-task conditions, separately for the early and late positivity windows. Because the late measurement window was selected from the target-minus-standard waveform, the overall late standard-versus-target comparison was treated as exploratory rather than an independent confirmatory test. All comparisons were performed on participant-level means using paired-samples tests.

For reliability analyses, two primary temporal ERP measures were retained: the N1-like amplitude averaged equally across stimulus types and task conditions, and the late target-minus-standard amplitude averaged equally across task conditions. Secondary analyses examined the constituent standard and target amplitudes, their differences, task-specific late oddball effects, and the early target related positivity. All ERP reliability measures used the same fixed measurement windows. One otherwise eligible temporal recording session was excluded because the 30-Hz-filtered ERP data required for quantification were unavailable.

#### Exploratory ERP decoding

As a complementary approach to conventional ERP amplitude analyses, we applied multivariate decoding to examine whether auditory responses contained information distinguishing standard from target tones. Given that the auditory oddball effect is well established in conventional EEG recordings, this analysis aimed to assess whether stimulus-specific information could also be recovered from the limited spatial coverage of the mobile EEG system by exploiting the temporal pattern of responses across electrodes. We performed time-resolved within-participant decoding of standard and target tones using session-averaged ERPs from the temporal electrodes (TP9 and TP10). Standard and target responses were averaged within each session across the two task conditions, with task conditions weighted equally. At each time point, a regularized logistic regression classifier used voltage samples from both temporal electrodes within a 40-ms window, advanced in 20-ms steps across the −200 to 800 ms epoch. Classification was evaluated using leave-one-session-out cross-validation, training on session-averaged responses from all but one session and testing on the held-out session. Accuracy was calculated separately for each participant and then averaged across the five participants. Statistical significance was assessed using 1,000 permutations that exchanged standard and target labels within sessions, preserving their paired structure. For each permutation, the classification analysis was repeated and the maximum group-level accuracy across the analysis window was retained to form a maximum-statistic null distribution. Corrected p-values were obtained by comparing observed accuracy at each window against this distribution, controlling family-wise error across time. This exploratory analysis tested generalization of stimulus-specific information across sessions, rather than classification of individual trials.

### Reliability analysis

We assessed the reliability of repeated EEG measurements from three complementary perspectives: within-person variability, between-person reliability, and the benefits of aggregating measurements across sessions. Analyses were performed for absolute measurement levels and, where applicable, within-session condition differences. First, we quantified within-person variability to characterize how much EEG measurements fluctuated across recording sessions within the same individual. For positive ratio-scale spectral measures, variability was expressed as the coefficient of variation (CV), which describes the standard deviation relative to the mean and allows comparisons across measures with different absolute magnitudes. For signed or non-ratio-scale measures, including ERP amplitudes and condition-difference scores, we used the standard deviation (SD), because the CV is not readily interpretable when measurements can approach or cross zero. Variability was calculated separately for each participant before being summarized across participants.

Second, we assessed between-person reliability to determine whether repeated recordings consistently captured individual differences in EEG measurements. For this purpose, we used a two-way, single-measure, absolute-agreement intraclass correlation coefficient (ICC[A,1]), which evaluates whether independently obtained estimates agree in their absolute values while accounting for variability both between and within participants. For each participant, two non-overlapping sets of recording sessions were randomly sampled from the eligible session pool, and measurements were averaged within each set to form two independent estimates. ICCs were then calculated across participants using these two estimates as measurement occasions. This procedure was repeated 200 times with a fixed random seed. Single-session reliability was estimated using one session per occasion, and aggregated reliability using four sessions per occasion. We report the mean ICC across repetitions and the 2.5th–97.5th percentiles of the resulting distribution. These intervals describe sensitivity to session selection rather than population-level confidence intervals.

Finally, we examined whether averaging repeated recordings improved the consistency of individual-level estimates. Although the ICC assesses agreement in absolute measurement levels, averaging multiple sessions may also improve how consistently participants can be distinguished from one another. To evaluate this, we repeated the random-split procedure for progressively larger sets of sessions. At each aggregation level, two non-overlapping sets of k sessions were sampled within each participant, and the Pearson correlation between the resulting participant-level estimates was calculated. Aggregation levels ranged from one session to the maximum number supported by the participant with the fewest eligible sessions, while retaining the same participants throughout each reliability curve. Correlations were averaged across 200 repetitions. Unlike the absolute-agreement ICC, these correlations quantify the consistency of between-participant differences rather than agreement in absolute measurement values. A correlation of .75 was included in figures as a descriptive reference and was not treated as a validated reliability threshold.

### Statistical analysis and software

All statistical analyses were performed in Python. Unless otherwise specified, statistical tests were two-sided with an alpha level of .05. EEG processing and analysis used MNE-Python (Gramfort et al., 2014), NumPy, SciPy, pandas, Pingouin (Vallat, 2018), scikit-learn (Pedregosa et al., 2011), and FOOOF/specparam (Donoghue, Haller, et al., 2020), with figures generated using Matplotlib. Randomized analyses, including permutation testing and session aggregation procedures, used fixed random seeds to ensure reproducibility.

## Results

### Spectral characteristics of the recorded EEG

Before assessing reliability, we first characterized the spectral properties of the EEG recordings across the eyes-open/eyes-closed and single-/dual-task conditions (Figure 1A). Power spectra at frontal (AF7, AF8) and temporal (TP9, TP10) electrodes showed the characteristic decrease in power with increasing frequency, with additional spectral structure superimposed on this aperiodic background. To distinguish periodic peaks from broadband activity, we parameterized the spectra using FOOOF (Donoghue, Haller, et al., 2020). The dashed curves in Figure 1A show the fitted aperiodic components, while the bars along the bottom of each panel indicate the participant-averaged proportion of sessions with a detected peak in each 1-Hz frequency bin. Peaks were frequently detected around 2–3 Hz, with additional peaks distributed across the theta, alpha, and beta ranges.

**Figure 1.**
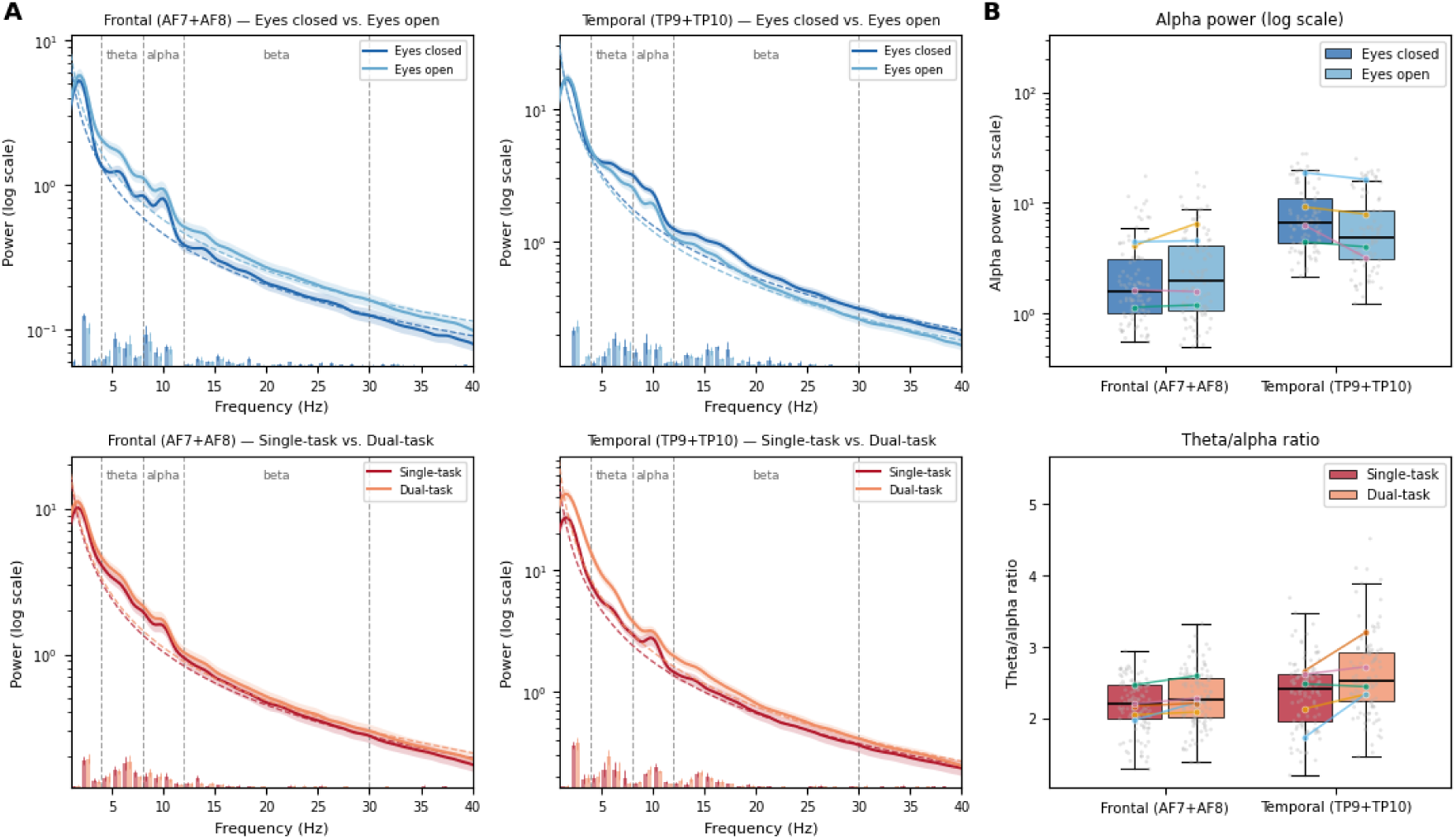
Spectral characteristics and sensitivity to experimental manipulations. (A) Mean ± SEM power spectra (log scale, 1–40 Hz) at frontal (AF7, AF8) and temporal (TP9, TP10) electrodes during eyes-closed and eyes-open rest (top row) and single- and dual-task oddball performance (bottom row). Dashed curves indicate the fitted aperiodic components, and dashed vertical lines mark conventional theta, alpha, and beta frequency-band boundaries. Short bars along the bottom of each panel indicate the participant-averaged proportion of sessions with a FOOOF-detected spectral peak within each 1-Hz frequency bin. (B) Distributions of alpha power during eyes-closed and eyes-open rest (top) and theta-to-alpha ratio during single- and dual-task performance (bottom), shown separately for frontal and temporal electrodes. Boxplots summarize session-level observations, with individual sessions shown as jittered points. Colored markers indicate participant-level means, with connecting lines linking each participant’s two conditions. Analyses included four participants for the eyes-open/eyes-closed contrast and five for the single-/dual-task contrast, following region-specific data-quality exclusions.

Alpha-range peaks (8–12 Hz) were detected in 48–83% of session-level spectra, depending on region and experimental condition. Their centre frequencies were relatively consistent across regions and conditions (mean 9.3–10.1 Hz), and all five participants exhibited alpha peaks in 50– 82% of their eligible session spectra when pooled across conditions, separately for each region. However, peak prevalence varied across recording conditions. At temporal electrodes, alpha peaks were more consistently detected than at frontal electrodes during task performance, where peak prevalence was approximately half of eligible session spectra. Frontal alpha peaks detected during task performance were also broader (mean bandwidth 3.3–4.1 Hz) than those detected during rest (2.0–2.2 Hz), while peak amplitudes above the fitted aperiodic background were relatively modest across regions and conditions. Although alpha-range peaks were detectable across all participants, their prevalence varied across recording sessions and conditions. These findings indicate that the Muse recordings contain identifiable spectral structure, while motivating a separate assessment of how consistently quantitative EEG measures can be estimated across repeated home recordings.

### Spectral sensitivity to experimental manipulations

We next examined whether spectral measures were sensitive to the experimental manipulations, comparing alpha power during eyes-open and eyes-closed rest and theta power, alpha power, and the theta-to-alpha ratio during single- and dual-task performance (Figure 1B). During rest, temporal alpha power was higher with eyes closed than eyes open in all four participants (mean difference = 1.81, 95% CI [−0.06, 3.68]), whereas frontal alpha power showed no consistent condition effect (mean difference = −0.61, 95% CI [−2.46, 1.24]). During the oddball task, theta power was higher under dual-task than single-task conditions in all five participants, both frontally (mean difference = 1.67, 95% CI [−0.21, 3.56]) and temporally (mean difference = 11.74, 95% CI [−10.93, 34.42]). Alpha power also tended to increase during dual-task performance, although this pattern was less consistent across participants (four of five frontally and three of five temporally). The theta-to-alpha ratio increased in all five participants at frontal electrodes (mean difference = 0.11, 95% CI [−0.00, 0.21]) and four of five at temporal electrodes (mean difference = 0.28, 95% CI [−0.06, 0.63]). Thus, despite substantial variation in effect magnitude, several spectral measures showed consistent directional sensitivity to the task manipulation across participants.

To examine whether these condition differences reflected frequency-specific activity or changes in the broadband spectral background, we compared the observed effects with those predicted by the fitted aperiodic components. Changes in the fitted aperiodic component predicted a substantial proportion of the mean task-related increases in theta power (49–52%) and particularly alpha power (70–103%). In contrast, only approximately 27–28% of the mean theta-to-alpha-ratio increase was accounted for by the aperiodic component. After subtracting the aperiodic-predicted changes, the frontal theta-to-alpha ratio difference remained positive in all five participants (mean residual = 0.078, 95% CI [−0.00, 0.16]), whereas the residual theta- and alpha-power effects were less consistent across individuals. During rest, the temporal alpha-power difference also remained positive after adjustment (mean residual = 1.14, 95% CI [0.08, 2.20]), indicating that this effect was not fully accounted for by changes in the aperiodic background. Together, these findings indicate that several spectral measures are sensitive to experimental manipulations, although the observed differences are partly associated with changes in the fitted aperiodic background. We next examined whether these condition-related spectral differences were sufficiently consistent across recording sessions to support within-participant decoding of experimental state.

### Cross-session decoding of experimental state

Having established that several spectral measures were sensitive to the experimental manipulations, we next examined whether these measures contained sufficient information to distinguish experimental conditions across recording sessions within individual participants. Using leave-one-session-out cross validation, a logistic regression classifier trained on 18 spectral features distinguished eyes-closed from eyes-open recordings with a mean accuracy of 76.2% across four participants (range: 50.0–88.0%; Figure 2A). This exceeded the group-level permutation distribution (95% interval: 42.2–57.1%, p = .005). Decoding exceeded the individual permutation distributions for three of the four participants, indicating that the group-level effect was not uniformly expressed across individuals. Classification of single- versus dual-task recordings was less accurate but also exceeded the group-level permutation distribution (mean accuracy: 63.7%, range: 46.0–84.4%; permutation 95% interval: 42.8–57.8%, p = .005). Although decoding accuracy was lower for the task contrast, performance varied considerably across participants, with four of five participants achieving accuracies above 50%.

**Figure 2.**
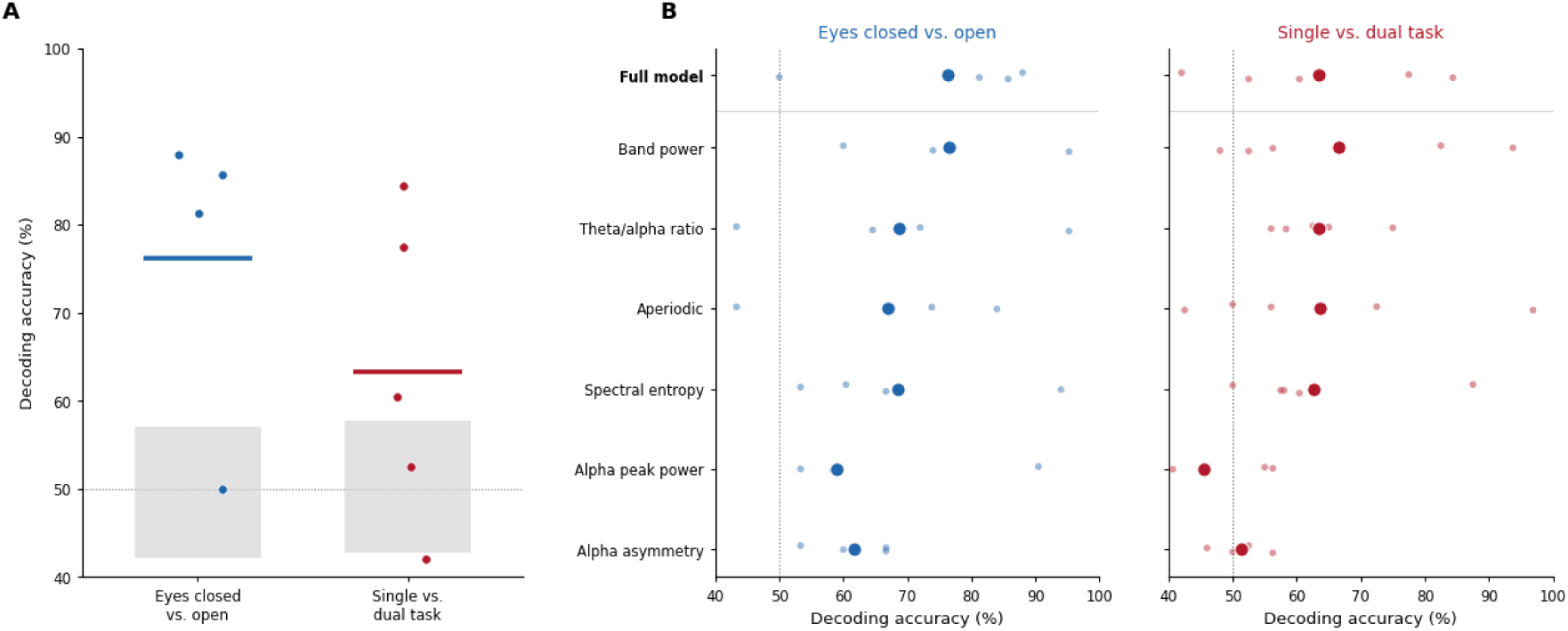
Cross-session decoding of experimental state from spectral EEG features. (A) Within-participant leave-one-session-out decoding accuracy for eyes-closed versus eyes-open rest (blue) and single- versus dual-task performance (red), using a logistic regression classifier trained on 18 spectral features. Small dots represent individual participants, and horizontal lines indicate group means. Grey shaded areas show the central 95% intervals of the group-level permutation-null distributions, obtained by randomly swapping condition labels within recording sessions (200 permutations). The dotted horizontal line indicates chance performance (50%). (B) Decoding accuracy using the full 18-feature model or six individual feature families, shown separately for the two experimental contrasts. Large dots indicate group means, and smaller, lighter dots represent individual participants. The dotted vertical lines indicate chance performance (50%). All models were evaluated using the same held-out recording sessions within each participant and contrast. Sessions failing quality control in either recording region were excluded from decoding. Analyses included four participants for the eyes-open/eyes-closed contrast and five for the single-/dual-task contrast.

To examine which spectral representations supported cross-session decoding, we repeated the analysis using six separate feature families (Figure 2B). For eyes-open versus eyes-closed recordings, band power (76.6%), the theta-to-alpha ratio (68.8%), aperiodic parameters (67.0%), and spectral entropy (68.6%) each supported substantial decoding accuracy when used independently, compared with 76.2% for the full model. A similar pattern emerged for the task contrast, where these four families achieved mean accuracies of 62.7–66.6%, close to the full model’s 63.7%. In contrast, alpha peak power and alpha asymmetry generally yielded lower decoding accuracies, particularly for the task contrast (45.6% and 51.4%, respectively). Nevertheless, the relative performance of individual feature families varied across participants. Together, these findings indicate that multiple spectral representations contain information about experimental conditions that generalizes across recording sessions. However, successful decoding does not necessarily imply that the underlying spectral measures are stable in absolute magnitude. We therefore next examined their reliability across repeated recordings.

### Reliability across repeated recordings

Having established that the recordings contained spectral information that generalized across sessions, we next examined the reliability of individual spectral measures across repeated recordings. Reliability was evaluated from three complementary perspectives (Figure 3). First, within-person variability quantified how much repeated measurements fluctuated within individuals. Second, intraclass correlations (ICCs) assessed how consistently individual differences in absolute spectral levels and condition-related effects were preserved across recording sessions. Finally, we examined whether combining information across increasing measures used to characterize the experimental manipulations above: the theta-to-alpha ratio during single- and dual-task performance and alpha power during eyes open and eyes-closed rest.

**Figure 3.**
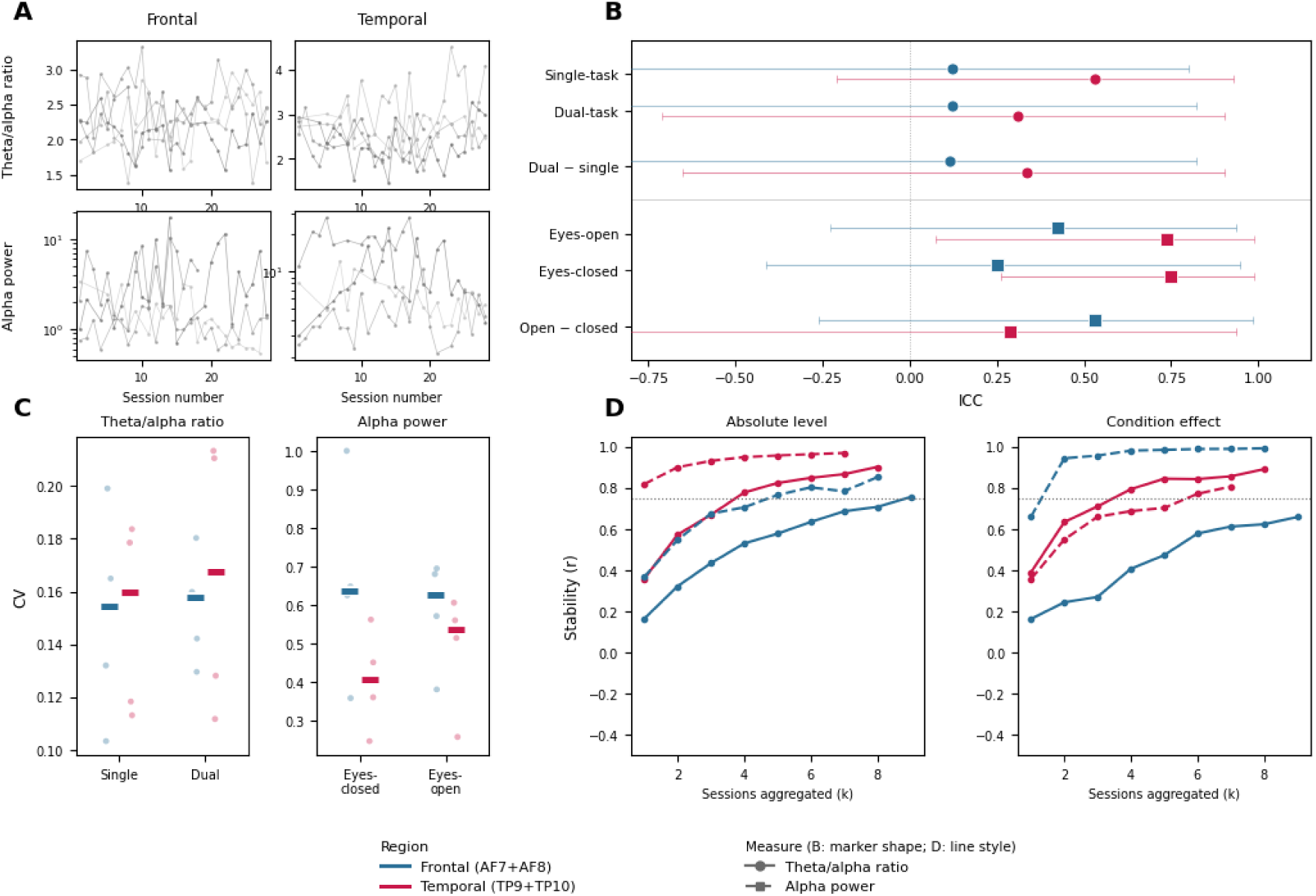
Session-to-session variability and reliability of spectral EEG measures. (A) Session-level trajectories of the theta-to-alpha ratio during single-task performance (top) and alpha power during eyes-closed rest (bottom), shown separately for frontal (AF7, AF8) and temporal (TP9, TP10) electrodes. Individual lines represent participants, illustrating variability across repeated recordings. Alpha power is plotted on a logarithmic scale. (B) Single-session absolute-agreement intraclass correlation coefficients (ICC(A,1)) for absolute spectral measurements and condition-related differences, shown separately for frontal (blue) and temporal (red) electrodes. Points indicate mean ICC estimates across 200 randomly sampled pairs of non-overlapping sessions, and horizontal lines indicate the 2.5th–97.5th percentile ranges across these samples, rather than population confidence intervals. Condition differences were calculated as dual minus single task for the theta-to-alpha ratio and eyes open minus eyes closed for alpha power. (C) Within-participant coefficients of variation (CVs) for the theta-to alpha ratio during single- and dual-task performance and alpha power during eyes-closed and eyes-open rest. Small dots represent individual participants, and horizontal bars indicate group means. (D) Split-half stability as a function of the number of sessions aggregated. For each aggregation level (k), two disjoint sets of k sessions were averaged within each participant, and Pearson correlations were calculated across participants between the resulting estimates. Curves show mean correlations across repeated random splits for absolute measurements (left) and condition-related differences (right), separately for frontal (blue) and temporal (red) electrodes. Solid lines indicate the theta-to-alpha ratio and dashed lines alpha power. Aggregation levels were restricted to those retaining a constant participant sample (up to nine frontal and eight temporal sessions per set for the theta-to-alpha ratio, and eight frontal and seven temporal sessions per set for alpha power). Analyses included five participants for the task contrast and four for the eyes-open/eyes-closed contrast. Sessions failing region-specific quality control were excluded only from the affected recording region.

The repeated-session trajectories revealed marked differences in within-person variability between measures (Figure 3A). Theta-to-alpha ratios remained relatively constrained across sessions, whereas alpha power showed substantially greater fluctuations. This difference was reflected in the within participant coefficients of variation (CVs; Figure 3C), which ranged from .15 to .17 for the theta-to alpha ratio across conditions and regions, compared with .41 to .66 for alpha power. Variability in alpha power was generally greater at frontal than temporal electrodes, whereas regional differences in theta-to-alpha variability were comparatively small.

Low within-person variability, however, does not necessarily imply reliable agreement across sessions. We therefore assessed the reliability of absolute spectral measurements using single-measure, absolute agreement ICCs estimated from randomly sampled pairs of recording sessions (Figure 3B). For the theta-to-alpha ratio, ICCs were low at frontal electrodes (.12 for both task conditions) but higher at temporal electrodes (.53 during single-task and .31 during dual-task performance). Alpha power showed a different regional pattern, with relatively high ICCs at temporal electrodes (.75 for eyes-closed and .74 for eyes-open rest), compared with lower values frontally (.25 and .42, respectively). Thus, despite its relatively low within-person variability, the frontal theta-to-alpha ratio showed limited single-session reliability, whereas temporal alpha power preserved individual differences more consistently across recordings.

We also examined the reliability of condition-related differences, calculated as dual minus single task for the theta-to-alpha ratio and eyes open minus eyes closed for alpha power. Condition-effect ICCs were low for frontal theta-to-alpha ratio (.11) and temporal alpha power (.29), but higher for temporal theta-to-alpha ratio (.34) and particularly frontal alpha power (.53; Figure 3B). Thus, condition-effect reliability varied considerably across measures and regions, with little evidence that subtracting conditions consistently improved reliability.

Although single-session reliability varied substantially across measures and regions, the present recording protocol was designed around relatively short but frequent EEG measurements, allowing information to be accumulated across sessions. We therefore examined whether averaging increasing numbers of recordings improved the stability of individual spectral estimates (Figure 3D). For absolute measurements, split-half correlations increased across all measures and regions, from .16–.82 when comparing individual sessions to .76–.97 when comparing averages of seven to nine sessions. Improvements were also evident for condition-related differences. Temporal theta-to-alpha-ratio condition effects showed a pronounced increase in stability (from .39 to .89), whereas frontal effects improved more modestly (from .16 to .66). For alpha power, temporal condition-effect correlations increased from .36 to .81, while frontal correlations were already relatively high for individual sessions (.66) and approached unity with aggregation (.99). However, these exceptionally high correlations should be interpreted cautiously given the small number of participants. Consistent with these findings, absolute-agreement ICCs increased for all 12 spectral measurements and condition effects when four sessions were aggregated, relative to a matched single-session baseline. Together, these results indicate that aggregating repeated short recordings can substantially improve the stability and absolute agreement of individual spectral estimates, although the magnitude of improvement varies across measures and regions.

### Auditory event-related responses and target-related activity

Whereas the preceding analyses focused on spectral characteristics of ongoing EEG activity, we next examined whether the mobile recordings also captured stimulus-locked auditory responses. Figure 4A shows the event-related responses to standard and target tones, collapsed across task conditions, separately for frontal and temporal electrodes. Event-related activity was substantially more pronounced at temporal electrodes, where both stimulus types elicited a positive-negative-positive sequence following tone onset. The most prominent feature was a negative deflection peaking approximately 375 ms after the logged stimulus onset (N1-like response; 325–425 ms measurement window), followed by a broader positive deflection extending into the late measurement window (591–691 ms). No comparable response morphology was apparent at the frontal electrodes. To determine whether the early auditory response differed according to stimulus type, we compared mean amplitudes elicited by standard and target tones within the N1-like window. This comparison revealed no significant difference (t(4) = −0.27, p = .802, dz = −0.12).

**Figure 4.**
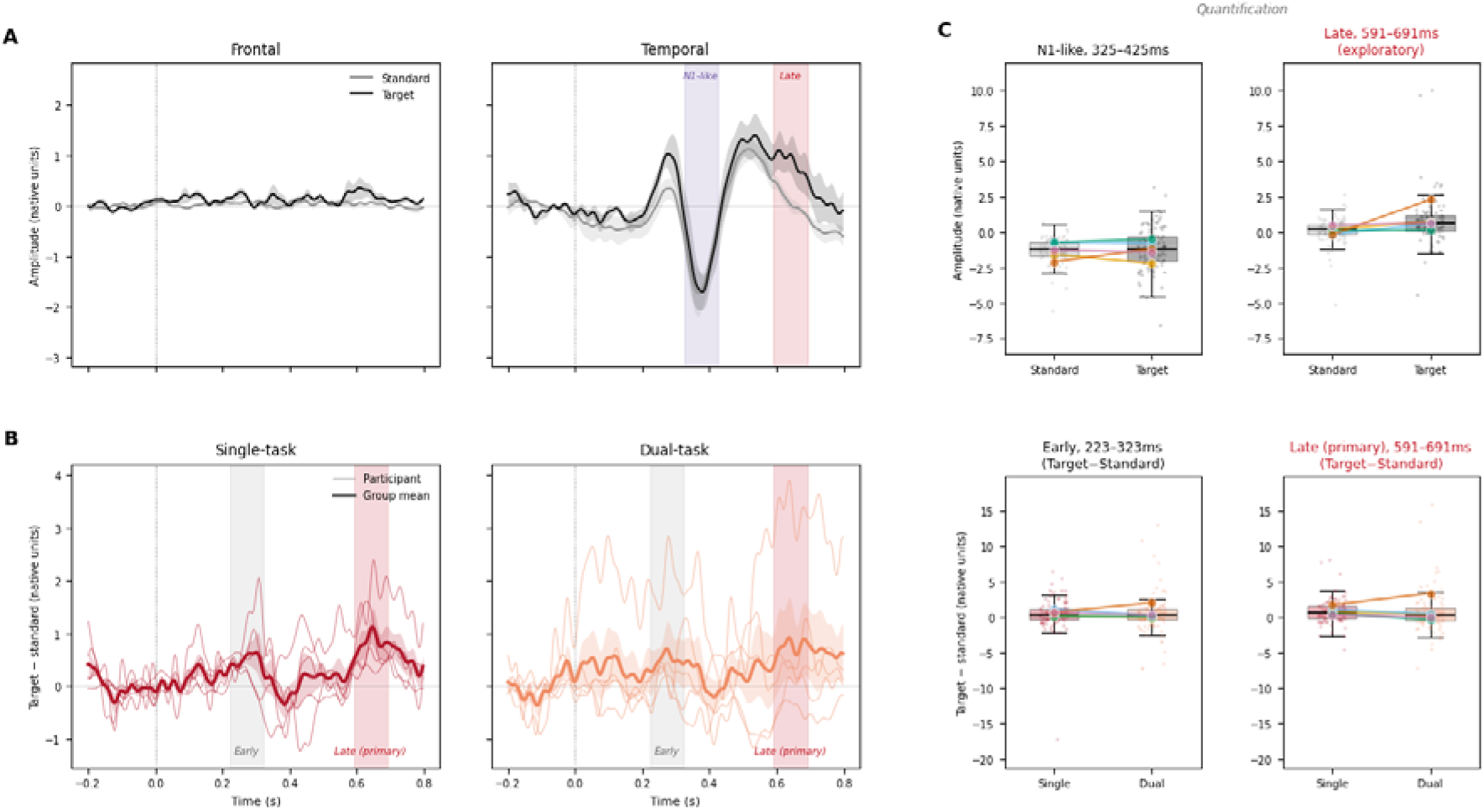
Auditory event-related responses and target-related activity during single- and dual-task conditions. (A) Grand-average event-related potentials (ERPs) recorded at frontal (AF7, AF8) and temporal (TP9, TP10) electrodes, separately for standard (gray) and target (black) tones. Shaded envelopes indicate variability across participants. Colored vertical bands mark the N1-like negativity (325–425 ms, purple) and late positivity (591–691 ms, red) measurement windows. (B) Temporal target-minus-standard difference waveforms during single-task (left) and dual-task (right) conditions. Thin lines represent individual participants, and thick lines indicate the group mean, with shaded envelopes indicating between-participant variability. Gray and red vertical bands indicate the early (223–323 ms) and late (591–691 ms) positivity windows, respectively. (C) Session-level amplitude distributions and participant-level means for the N1-like response (top left), exploratory late standard versus-target comparison (top right), and early (bottom left) and late (bottom right) target-minus-standard responses during single- and dual-task conditions. Boxplots summarize session-level distributions, light points represent individual sessions, and colored connected markers represent participant means. All amplitudes are expressed in native device units. Vertical dotted lines in panels A and B indicate logged stimulus onset (0 ms). Analyses included five participants (P02–P06) and 105 eligible paired temporal recording sessions.

Whereas the N1-like response primarily characterized the stimulus-locked auditory activity, we next examined whether the recordings captured additional activity associated with target detection. Target-minus-standard difference waveforms showed an early positive deflection (223– 323 ms) and a more pronounced late positivity (591–691 ms; Figure 4B). In the late window, target amplitudes were numerically larger than standard amplitudes in all five participants (mean target-minus-standard difference = 0.80 native units, 95% CI [−0.45, 2.04]). However, the comparison was not statistically significant (t(4) = 1.78, p = .150, dz = 0.80). Because the late window was identified from the target-minus-standard waveform itself, this comparison was treated as exploratory rather than an independent confirmatory test.

Finally, we examined whether target-related activity differed between single- and dual-task conditions (Figure 4C). Neither the early positivity (mean dual-minus-single difference = 0.01, 95% CI [−1.01, 1.03], t(4) = 0.03, p = .976, dz = 0.01) nor the late positivity (mean difference = 0.09, 95% CI [−1.16, 1.35], t(4) = 0.20, p = .851, dz = 0.09) showed a consistent task-load modulation. Thus, although temporal recordings captured clear stimulus-locked auditory activity and a descriptively positive late target-related response, the present sample did not establish reliable modulation of these ERP amplitudes by cognitive load.

### Cross-session decoding of auditory stimulus identity

Although the late target-related positivity was descriptively present in all five participants, the conventional mean-amplitude analysis did not establish a statistically reliable difference between targets and standards. We therefore explored whether stimulus-specific information could be detected using a complementary multivariate approach that considered patterns of activity across temporal electrodes and neighboring time points. Specifically, we performed time-resolved decoding of standard and target tones using session-averaged ERPs from the temporal electrodes, with leave-one-session-out cross-validation to assess generalization across recording sessions (Figure 5). Decoding accuracy increased above chance approximately 200 ms after tone onset and remained elevated throughout much of the subsequent analysis window. Statistically significant decoding was observed at 300–340 ms and 420–480 ms (corrected ps = .010–.014), as well as 480–520 ms (corrected p = .041), following maximum-statistic correction across the analysis window. Peak accuracy reached 64.3% at 440–480 ms, with all five participants showing numerically above-chance performance at this interval (55.3–75.0%). No prestimulus window showed significant decoding after correction. Thus, despite the absence of a statistically reliable target-related mean-amplitude difference, the temporal ERP waveforms contained information distinguishing standards from targets that generalized across recording sessions.

**Figure 5.**
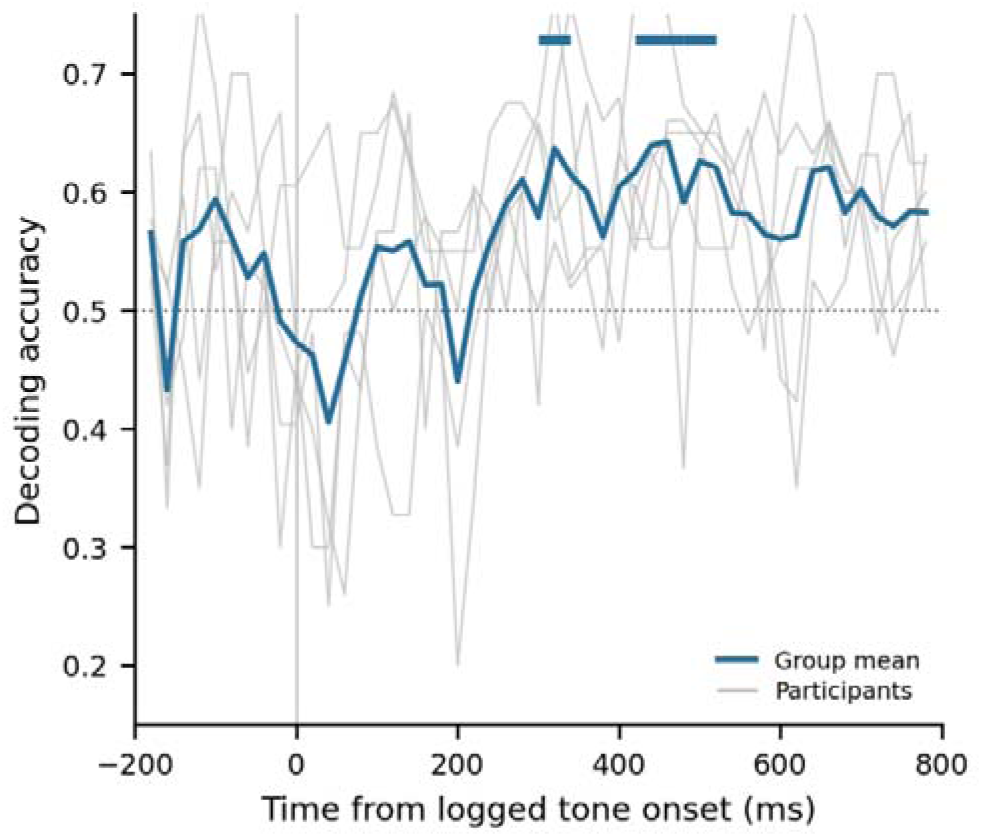
Exploratory time-resolved decoding of auditory stimulus identity across recording sessions. Decoding accuracy for distinguishing standard and target tones from session-averaged event-related potentials recorded at temporal electrodes (TP9 and TP10). Classification was performed separately within each participant using leave-one-session-out cross-validation. The thick blue line shows mean classification accuracy across five participants (105 recording sessions), and thin grey lines show individual-participant accuracy. The horizontal dotted line indicates chance performance (50%), and the vertical grey line marks logged tone onset (0 ms). Blue horizontal bars indicate individual 40-ms windows with above-chance decoding after family-wise error correction across the full analysis interval using a maximum-statistic permutation procedure (1,000 permutations). Sliding windows were advanced in 20-ms steps. The analysis was exploratory and evaluated classification of session averaged responses rather than individual trials.

### Reliability of auditory event-related responses

We next examined whether the auditory ERP responses identified above could be measured reliably across repeated recording sessions. We focused on two measures derived from the temporal electrodes: the stimulus-collapsed N1-like response (325–425 ms) and the late target-minus-standard positivity (591–691 ms). Across five participants and 105 eligible sessions, both measures showed substantial session-to-session variability, particularly the late target-related response (mean within-person SD = 1.39 versus 0.83 native units for the N1-like response; Figure 6A).

**Figure 6.**
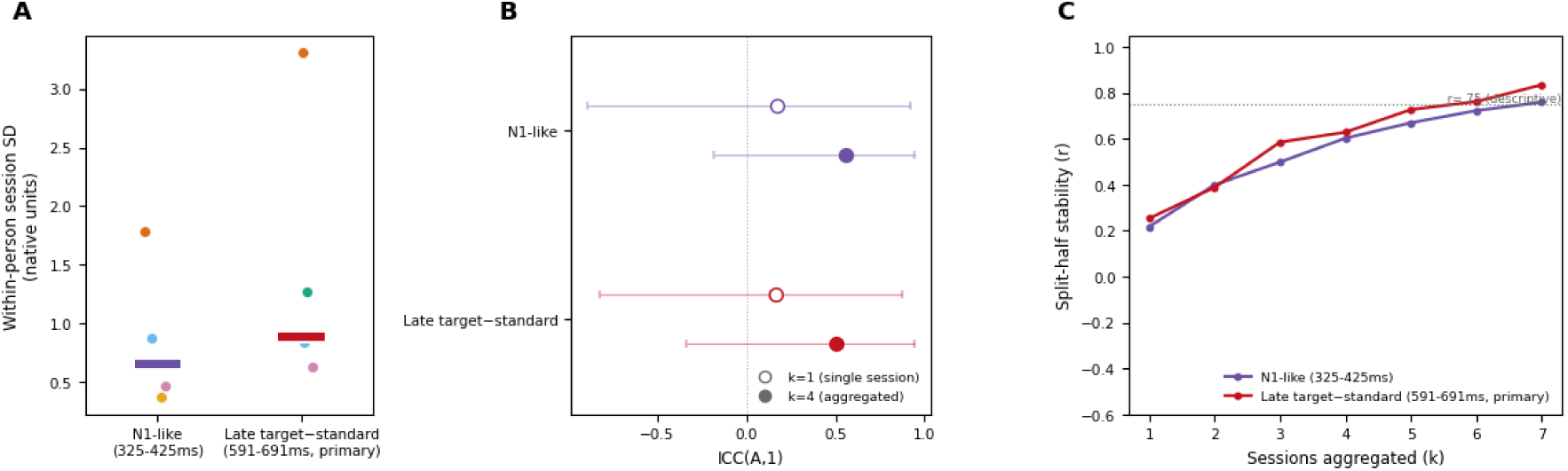
Reliability of auditory event-related potential measurements across repeated recording sessions. (A) Within-participant standard deviation (SD) of session-level amplitudes for the stimulus-collapsed N1-like response (325–425 ms) and late target-minus-standard positivity (591–691 ms), both measured at temporal electrodes. Colored points represent individual participants, and horizontal bars indicate group medians. (B) Single-session (open circles) and four-session-aggregated (filled circles) absolute-agreement intraclass correlations (ICC[A,1]) for the same measures. Points indicate mean ICC estimates, and horizontal intervals indicate the 2.5th–97.5th percentiles across 200 random selections of non-overlapping session sets. These intervals reflect variability due to session selection rather than population-level confidence intervals. (C) Split-half Pearson correlations across participants as a function of the number of sessions averaged within each half. Curves represent mean correlations across 200 random splits, with a maximum of seven sessions per half. The horizontal dotted line at r = .75 is included as a descriptive reference, not a validated reliability threshold. Analyses included five participants and 105 eligible temporal recording sessions; the same participants contributed to every aggregation level.

To determine whether averaging repeated recordings improved reliability, we compared estimates based on individual sessions with estimates obtained by averaging measurements across four sessions per participant. Single-session absolute-agreement ICCs were modest for both the N1-like response (ICC = .17) and the late target-related positivity (ICC = .17; Figure 6B). Averaging four sessions per estimate increased ICCs to .56 and .50, respectively, although estimates varied considerably across random session selections. This improvement was also apparent when examining split-half consistency across progressively larger sets of sessions (Figure 6C). Correlations increased from r = .22 to .76 for the N1-like response and from r = .25 to .83 for the late target-minus-standard positivity as the number of sessions increased from one to seven per half. Together, these findings suggest that aggregating repeated recordings can substantially improve the consistency of individual-level ERP measurements. However, given the small number sensitive to individual differences in recording quality and between-participant variability.

Finally, we examined whether the benefits of aggregation extended to other ERP measures. Reliability differed substantially depending on the component and contrast of interest. Within the N1-like window, standard-tone amplitudes were more reliable than target amplitudes or their difference (split-half r at seven sessions: .86, .57, and .36, respectively). Similarly, the late target-minus-standard positivity showed greater reliability during dual-task than single-task performance (r = .79 versus .28), despite the absence of a consistent task-load effect on mean amplitude. The dual-minus-single difference itself showed relatively low reliability (r = .37). These findings indicate that, although repeated recordings can improve the reliability of ERP measurements, this benefit does not extend equally to all measures, particularly contrasts intended to capture individual differences in cognitive-load effects.

## Discussion

Mobile EEG offers considerable promise for studying brain activity beyond conventional laboratory settings, enabling repeated measurements across days under more naturalistic conditions (Sabio et al., 2024). Previous studies have demonstrated that wearable EEG systems can capture meaningful neural signals outside the laboratory, including through repeated, self-administered recordings (Barbey et al., 2022; Mikkelsen et al., 2021). However, the reliability of spectral and event-related measures obtained using low-density consumer EEG across repeated recording sessions remains less well established. Here, we addressed this question by combining brief, repeated at-home recordings with controlled resting-state and auditory oddball paradigms. Our findings demonstrate that mobile EEG recordings contain meaningful information about both physiological state and task demands. Spectral features distinguished eyes-open from eyes-closed resting states and single-from dual-task performance, with classification generalizing across recording sessions. Moreover, we observed clear stimulus-locked auditory responses at the temporal electrodes, alongside a descriptively positive late target-related response. Together, these findings support the feasibility of using mobile EEG to capture both ongoing oscillatory activity and event-related responses across repeated recordings outside the laboratory.

Beyond demonstrating the presence of meaningful electrophysiological signals, an important question is whether these signals can be measured reliably across recording sessions. This is particularly relevant for mobile EEG, where recording conditions may vary from day to day and individual sessions are often relatively short. Previous research has demonstrated encouraging test–retest reliability for spectral measures obtained with consumer EEG devices (Rogers et al., 2016), while repeated home recordings using more extensive wearable systems have shown that aggregating measurements can improve ERP precision (Barbey et al., 2022). Extending these findings to a low-density consumer EEG system, our results revealed considerable session-to-session variability in both spectral and event-related measures, with single-session reliability differing substantially across features. Notably, however, aggregating information across repeated sessions improved the stability of individual-level estimates. This improvement was evident for several spectral measures as well as auditory ERP amplitudes, suggesting that repeated sampling can help overcome some of the limitations associated with individual recordings. Importantly, these findings should not be taken to imply that individual mobile EEG sessions are inherently unreliable or that aggregation across days is always necessary. The recordings used here were relatively brief, and measurement reliability is likely to depend on the amount of usable data collected within each session, as well as the feature being measured. Aggregating sessions is particularly useful when the objective is to obtain stable estimates of individual characteristics. In contrast, studies seeking to track changes in neural activity over time must balance measurement precision against temporal resolution, as averaging across extended periods may obscure meaningful fluctuations.

Importantly, the spectral differences observed across experimental conditions were broadly consistent with established electrophysiological signatures of changes in physiological state and cognitive demands. Eye closure was associated with increased temporal alpha power, in line with the well established modulation of alpha activity by visual input (Adrian & Matthews, 1934; Barry et al., 2007). Similarly, increasing cognitive load through concurrent mental arithmetic was accompanied by increases in theta power and the theta-to-alpha ratio, consistent with previous research linking spectral EEG activity to cognitive workload (Chikhi et al., 2022; Gevins et al., 1997). However, our findings also highlight the importance of how these spectral changes are quantified. A substantial proportion of the observed increases in theta and alpha power during dual-task performance could be explained by changes in the fitted aperiodic spectral background, whereas the theta-to-alpha ratio was considerably less affected. Moreover, the theta-to-alpha ratio showed substantially lower relative session-to-session variability than alpha power, consistent with recent evidence that spectral power ratios can provide relatively stable measurements across recording sessions (Chang, 2025). Together, these findings suggest that spectral features derived from repeated mobile EEG recordings can distinguish between different levels of cognitive demand, with the theta-to-alpha ratio emerging as a particularly informative measure due to its sensitivity to task-related changes and relatively low within-person variability. Nevertheless, because spectral ratios can reflect multiple periodic and aperiodic contributions (Donoghue, Dominguez, et al., 2020), their physiological interpretation requires caution.

The event-related analyses further demonstrate the potential of low-density mobile EEG to capture meaningful neural responses during repeated at-home recordings. We observed a clear stimulus-locked auditory response at the temporal electrodes, consistent with previous demonstrations that consumer grade EEG systems can measure auditory event-related activity (Krigolson et al., 2017). Although a late target-related positivity was descriptively present across participants, the corresponding mean amplitude difference did not reach statistical significance. Nevertheless, exploratory time-resolved decoding revealed that standards and targets could be distinguished from their session-averaged ERP waveforms, with classification generalizing across recording sessions. This finding is consistent with previous research demonstrating that multivariate approaches can detect stimulus-related information that may be difficult to establish using conventional ERP amplitude comparisons (Carrasco et al., 2024; Grootswagers et al., 2017). Importantly, repeated recordings also allowed substantial numbers of auditory trials to be accumulated across days, potentially improving the precision of individual ERP estimates. Consistent with this possibility, aggregating sessions progressively improved the stability of both the condition-general auditory response and the late target-related positivity. These findings highlight the potential of repeated mobile EEG measurements to improve the stability of event-related estimates by distributing data collection across multiple shorter recordings rather than relying on a single prolonged session.

Several considerations should be taken into account when interpreting these findings. First, although the repeated-measures design provided extensive recordings within individuals, the number of participants was relatively small. Consequently, estimates of between-person reliability and the generalizability of the observed effects should be interpreted cautiously, and future studies with larger samples will be important for establishing how consistently these findings extend across individuals. Second, the four-channel EEG system provides limited spatial coverage, and its common frontal reference complicates the interpretation of regional differences in the recorded signals. Accordingly, although certain responses were more pronounced at temporal than frontal electrodes, these differences should not be interpreted as evidence of their anatomical origin. Finally, self-administered home recordings introduce variability in electrode contact and recording conditions, particularly when participants independently reposition the headband at the start of each session. Nevertheless, meaningful electrophysiological measurements could be obtained despite repeated repositioning. Consumer EEG headbands can be fitted with relatively little training and without the extensive electrode preparation required for conventional EEG systems. Combined with their relatively low cost and accessibility, this makes repeated data collection practical. An important direction for future research is therefore to establish how recording duration, number of usable trials, and sampling frequency jointly determine the reliability of different mobile EEG measures. Longer individual recordings may improve measurement precision, whereas repeated shorter sessions may offer advantages in terms of feasibility and participant adherence. The optimal balance will depend on the intended application, particularly whether the goal is to characterize stable individual differences or detect changes within individuals over time. Establishing the magnitude of measurement variability relative to meaningful longitudinal changes will be essential for determining whether mobile EEG can reliably capture changes in neural activity across days, weeks, or longer periods.

In conclusion, the present study demonstrates the feasibility of obtaining meaningful spectral and event-related EEG measurements through repeated, self-administered recordings using a low-density consumer device. Across resting-state and cognitive-task conditions, the recordings captured established electrophysiological signatures of physiological state and cognitive demands, as well as stimulus-specific auditory responses. Although measurement stability varied across features and recording sessions, our findings demonstrate that repeated sampling can improve the precision of individual-level estimates. More broadly, this work provides a foundation for developing reliable longitudinal EEG measurements outside the laboratory. By optimizing recording duration, sampling frequency, and analytical approaches for different research questions, mobile EEG may ultimately enable researchers to characterize not only stable individual differences but also meaningful changes in neural activity over time. Beyond longitudinal home monitoring, the accessibility and portability of consumer EEG systems may facilitate research in more naturalistic settings, including driving, virtual reality, and gaming, as well as future applications in cognitive-state monitoring and digital health.

